# Detailed microbiome analysis of sticker-stripped surface materials of acne lesions revealed acne-related *Cutibacterium acnes* subtypes: a pilot study

**DOI:** 10.1101/2023.12.19.569832

**Authors:** Yutaka Shimokawa, Osamu Funatsu, Kazuma Ohata, Fukashi Inoue, Kota Tachibana, Itaru Dekio

## Abstract

*Cutibacterium acnes (C. acnes)* is known to play a central role in pathogenesis of acne vulgaris. It has been understood that multiple phylotypes of *C. acnes* exist, with certain types being more prevalent in patient with acne vulgaris and others more common in healthy individuals. In this context, we conducted a preliminary study using self-collected samples via an adhesive sticker (MySkin® patch) to analyze the skin microbiome of Japanese women. The study aimed to determine the role of *C. acnes* and its specific phylotypes in the development of acne vulgaris.

Participants in this study were Japanese females aged between their 20s and 40s. Dermatologists evaluate the data from web-based questionnaires and smartphone image submissions to classify subjects into either Acne group (n = 219) or Non-acne group (n = 77). Quality assessment of DNA extracted from the sticker was conducted, followed by amplification of the *16S rRNA* region using PCR. Subsequent microbial community analysis was performed using next-generation sequencing techniques. Genetic classification of *C. acnes* was accomplished through single locus sequence typing.

Results indicated a bacterial community composition on the facial skin surface predominantly consisting of *C. acnes* clusters, with over half of these clusters constituted by *C. acnes*. Notably, the Acne group exhibited a significantly higher proportion of *C. acnes* relative to total bacterial presence compared to the Non-acne group. Analysis of *C. acnes* phylotypes revealed a markedly lower presence of type III (subspecies *elongatum*) in the Acne group (vs. Non-acne group, *p* < 0.05). No significant differences were observed in the prevalence of Types IA_1_, IA_2_, II, and IB between the two groups. The predominantsequence types (ST) of *C. acnes* identified were IA2_2_F0 (23.9%), IA1_4_A0 (20.6%), and II_2_K0 (18.6%). Within the Acne group, an increase in IA2_1_F1 and a decrease in III_1_L0 were observed (vs. Non-acne group, *p* < 0.05).

This study underscores the feasibility of using self-collected and mailed-in samples for qPCR and microbiome analysis, maintaining diagnostic quality comparable to in-person assessments. Furthermore, the variation in the expression of *C. acnes* phylotypes across skin surfaces between acne-afflicted and healthy individuals could suggest that shifts in phylotype expression patterns may be indicative of skin susceptibilities to acne development.

## Introduction

Human skin serves as a barrier that separates the external and internal environments. It prevents the invasion of harmful substances into the body and blocks the loss of moisture and beneficial substances from the body. Its surface is home to many microorganisms, including bacteria, fungi, and viruses, which form colonies. These microorganisms interact with each other and with their host, creating a stable and complex ecosystem through cross-talk. This skin microbiome changes with factors such as age, gender, temperature, pH, ultraviolet exposure, moisture and humidity, and sebum content^1,2^. In recent years, comprehensive analyses of the skin microbiome have been conducted using next-generation sequencing techniques, specifically the analysis of bacterial *16S rRNA* gene. Reports have been emerging about its involvement in inflammatory skin diseases such as acne vulgaris, atopic dermatitis, and psoriasis^3–6^.

Among the skin microbiome, the most common commensal bacterial species is *Cutibacterium acnes*^7,8^. *C. acnes* is a ubiquitous Gram-positive anaerobic bacterium belonging to the phylum *Actinobacteria* and is primarily found deep within the sebaceous hair follicles where it comes into contact with keratinocytes^9^. Furthermore, *C. acnes* is known to metabolize free fatty acids from sebum, which reduces the surface pH. This acidic environment potentially inhibits pathogenic bacteria such as *Staphylococcus aureus* and promotes the growth of other healthy commensal bacteria, suggesting its contribution to barrier homeostasis^10–12^.

Based on genotyping, it has become clear that *C. acnes* has multiple subtypes, ranging from those involved in the onset of acne vulgaris to those existing on healthy skin^13–15^. Moreover, the subspecies of *C. acnes* are classified as *C. acnes* subspecies *acnes* (*C. acnes* type I), subspecies *defendens* (*C. acnes* type II), and subspecies *elongatum* (*C. acnes* type III)^16,17^. Further, based on single locus sequence typing (SLST) and whole genome sequencing (WGS) methods, the genetic sequence analysis of this species has been phylogenetically refined and categorized into subtypes IA_1_, IA_2_, IB, IC, II, and III^17^. Among these, subtype IA_1_ of *C. acnes* has been isolated from acne vulgaris and is believed to have strong pathogenicity^13,18,19^. On the other hand, subtypes IB^15^ and II^14^ are expressed in healthy skin and are not believed to be associated with the onset of acne vulgaris^19^. Thus, while there are multiple subtypes of *C. acnes* that seem to have different roles, many aspects of their detailed ecology remain unknown.

At present, various methods such as cotton swabs, scraping, cyanoacrylate gel biopsy, and needle biopsy are used to collect skin bacteria for analysis of the skin microbiome. However, reports suggest that the detection rate of *C. acnes* varies depending on the sampling method^20–22^. The most common sampling method is the swabbing method using a cotton swab^23,24^, but it is difficult to control the sampling efficiency depending on the collection site and technique, and it has the disadvantage of poor preservation^20^. In any case, it has not been envisaged that subjects would collect their own samples, and until now, it has been difficult to collect samples by mail and analyze the skin microbiome. However, with recent technological advancements, a sticker stripping kit for self-sampling called MySkin® patch, which possesses features like preservation and quantification and can withstand outdoor temperatures for postal services, has been developed. Therefore, this study conducted a preliminary investigation with the aim to elucidate the relationship between the skin resident bacterium *C. acnes* and acne vulgaris using samples collected by self-sampling with the sticker stripping kit MySkin® patch among Japanese women.

## Materials and Methods

### Samples

From the female Japanese customers in their 20s to 40s who had purchased products from KINS Co., Ltd., 296 individuals who agreed to the content of this study were selected as the subjects. This research was conducted based on the Declaration of Helsinki and the ethical guidelines for medical research, with the approval by the SOUKEN Ethics Committee in April 2021 (Approval number: 146853_rn-30080). Individuals who had used antibiotics within a month prior to sample collection were excluded from the study. Subjects were not given any lifestyle restrictions such as the use of skincare products, except just before sampling. The condition of the facial skin was determined from a web questionnaire and a smartphone photo image of the face. The web questionnaire acquired responses to the question, “Do you currently have visible pimples or acne on your face? Yes/No,” and concurrently, subjects sent photos taken with their smartphones. Dermatologists evaluated these photos and categorized the patients into Acne group/Non-acne group. Subjects, after waking up and before washing their faces, applied the dedicated adhesive sticker MySkin® patch (an oval-shaped sticker with the area of 6.4 mm^2^) to the specified locations on their facial skin for 1 min to collect samples. After collection, the samples were promptly mailed to the analytical institution. The flow from sample collection to data analysis is shown in Fig. 1. The dedicated skin sticker used in this study, designed for collecting skin microbes, has superior microbial preservation and has been demonstrated to monitor relative bacterial count fluctuations^25^.

**Figure 1.**
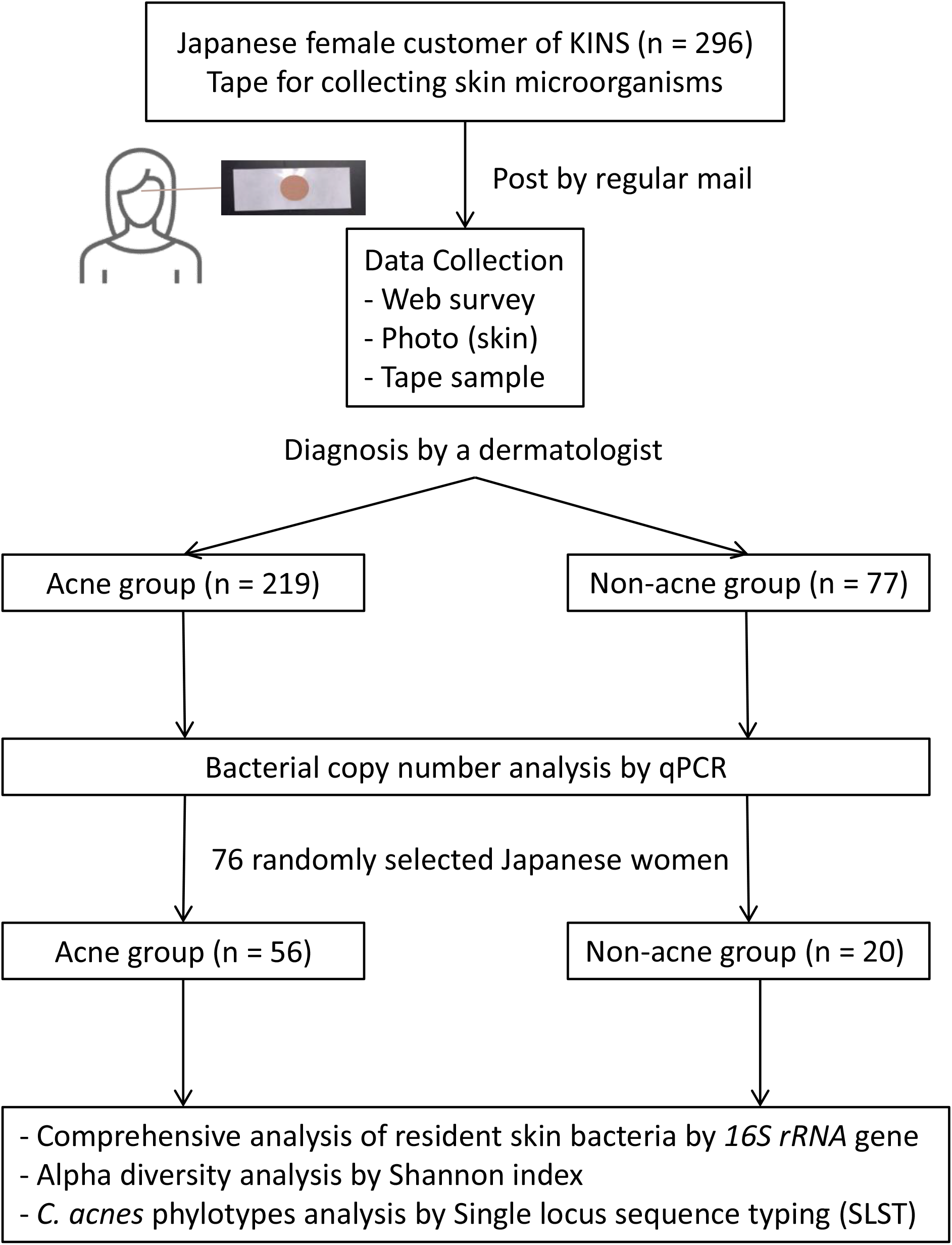
A simplified schema summarizing the skin microbiome analysis. Skin sampling was conducted using the sticker-stripping method (for 1 minute, before face wash in the morning).

### Skin microbiome analysis

Bacterial community analysis was performed by MySkin Co., Ltd. The DNA was extracted from the skin sticker fragments and after performing a Quality Check (QC), the obtained DNA was used as a template to amplify the bacterial *16S rRNA* by PCR. Subsequently, indexes for next-generation sequencer migration were added to the PCR. The DNA concentration of each sample was measured, and all samples were adjusted to be of the same concentration. The adjusted DNA was purified and then sequenced using a next-generation sequencer (MiSeq, Illumina Inc.) to produce the library. The library was analyzed using the next-generation sequencer, targeting the V1-V2 regions of the *16S rRNA* gene for bacterial community analysis. The microbiome analysis tool QIIME2 was used, and for bacterial species prediction, the *16S rRNA* gene database Greengenes was employed. For the single gene-based subtype structure analysis of *C. acnes*, a *C. acnes*-specific genomic region that can classify into six subtypes (IA_1_, IA_2_, IB, IC, II, III) was utilized. A unique database regarding *C. acnes* was used to determine the subtypes. Bacterial species in each sample were summarized by taxonomic rank, and from the number of 16S rRNA reads, the relative abundance of each bacterium was calculated, and the bacterial composition and percentages were determined. Based on these data, hierarchical clustering using the Ward’s method was performed to group the microbial communities. Alpha diversity was evaluated using the Shannon index^26^. Moreover, for the *C. acnes* subtype analysis, in conjunction with the *16S rRNA* gene-based microbiome analysis, a single locus sequence typing (SLST) analysis of *C. acnes* was conducted.

Polymerase chain reaction (PCR) was conducted using ExTaq (TaKaRa) to amplify the SLST locus. Following amplification, the PCR products were purified using magnetic beads (Sera-Mag™ Select, Cytiva). These purified samples were then analyzed using the MiSeq sequencing platform. The resulting sequences were trimmed and classified into SLST classes using the self-built SLST database.

The primer sequences used for the SLST analysis are shown below.

Fw-R3B:TCGTCGGCAGCGTCAGATGTGTATAAGAGACAGacaccagggggtcaacttgg

Rv-R3AB:GTCTCGTGGGCTCGGAGATGTGTATAAGAGACAGatgttacccatgtaatgggcagg

The uppercase represents the adapter sequences for MiSeq sequencing, while the lowercase indicates the sequences used for acne bacteria typing.

### Copy number analysis by qPCR

For *C. acnes*, the copy numbers in the materials were determined by calculating bacterial cell numbers per fixed area of the sticker from qPCR quantification. First, the skin adhesive sticker fragments were placed in a tube and immersed in 1 ml of buffer, followed by a spin-down and then left at 100 ℃ for 15 minutes. For the residual liquid after removing the sticker, nucleic acid extraction was performed using the phenol/chloroform method. A standard solution was adjusted and used by stepwise dilution of the standard DNA with a known number of copies. The sequence of the PCR primers is confidential, but it is possible to provide information upon request. PowerUp™ SYBR™ was used for detection, and nucleic acid amplification was performed using the qPCR device (QuantStudio, Applied Biosystems).

### Statistical Analysis

The measurement results are shown as mean ± standard error, and if the *p*-value is less than 0.05, it was judged as statistically significant. For comparisons between two groups, Welch’s *t*-test was used, and for comparisons among three or more groups, one-way analysis of variance (ANOVA) and *post-hoc* testing using the Bonferroni method were appropriately applied. Statistical processing was performed using HAD^27^.

## Results

### Sample Quality

Quality control of the DNA extracted from the mailed samples was conducted. After PCR (both the first and second times), agarose gel electrophoresis was performed to confirm that the PCR product was of the expected size (data not shown). In addition, quantification was carried out using qPCR, utilizing PCR primers corresponding to a portion of the P5 and P7 sequences to verify that adapter sequences had been added to both ends of the library. Based on these verifications, it was determined that the extracted DNA was suitable for use in the *16S rRNA* gene-based microbiome analysis.

### Skin microbiome analysis

Out of 296 participants aged between 20s and 40s (Table 1), a comprehensive analysis of the resident bacteria on the facial skin was initially conducted on 76 randomly selected Japanese women (56 in the Acne group and 20 in the Non-acne group). As a result of hierarchical clustering analysis, the bacterial flora was classified into three groups: *C. acnes*-dominant, *Neisseriaceae*-dominant, and others (Fig. 2-A). The most prevalent was the cluster where *C. acnes* was dominant, in which 50-90% or more was composed of *C. acnes*. Subsequently, there were many clusters dominant with *Neisseriaceae*, which are resident bacteria of the oral cavity. On the other hand, regarding the fungal flora, it was divided into five major groups, consisting of clusters dominated by *Malassezia restricta* and *Malassezia globosa*, which are also suggested to be involved in acne vulgaris, and other clusters (Fig. 2-B). Interestingly, in the other clusters, there were participants who were dominated by *Rothia mucilaginosa*^28^, which is a resident bacterium of the oral cavity, and *Wickerhamomyces anomalus*^29^, known as a pathogenic yeast.

**Figure 2.**
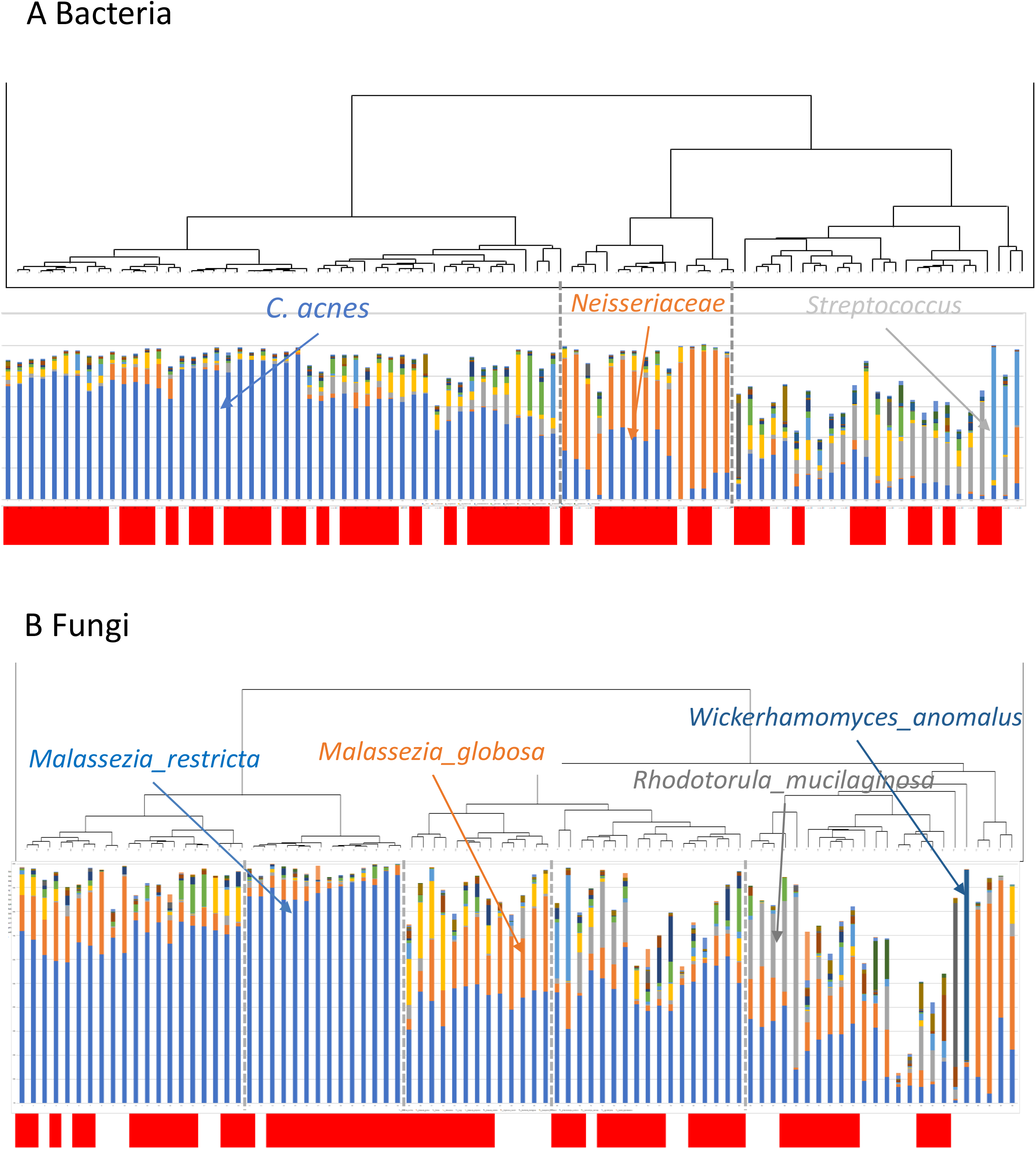
Taxonomic composition of bacteria (A) and fungi (B) based on *16S rRNA* and ITS sequences. Acneic subjects are highlighted in red bar at the bottom of figure.

**Table 1.** Subject characteristics.

| Characteristics | Acne | Non-acne |
| --- | --- | --- |
| n | 219 | 77 |
| Age (average) | 40.3 | 34.7 |

### Alpha diversity

The ***α***-diversity (species richness) on the facial skin was compared between the bacterial and fungal communities. The Shannon index for bacterial communities showed a median value of 1.35 in the Acne group and 1.66 in the Non-acne group, with the Acne group being significantly lower (*p* < 0.05, Fig. 3-A). On the other hand, there was no noticeable difference in the α-diversity of the fungal communities between the two groups (Fig. 3-B). Therefore, subsequent analyses focused solely on bacteria.

**Figure 3.**
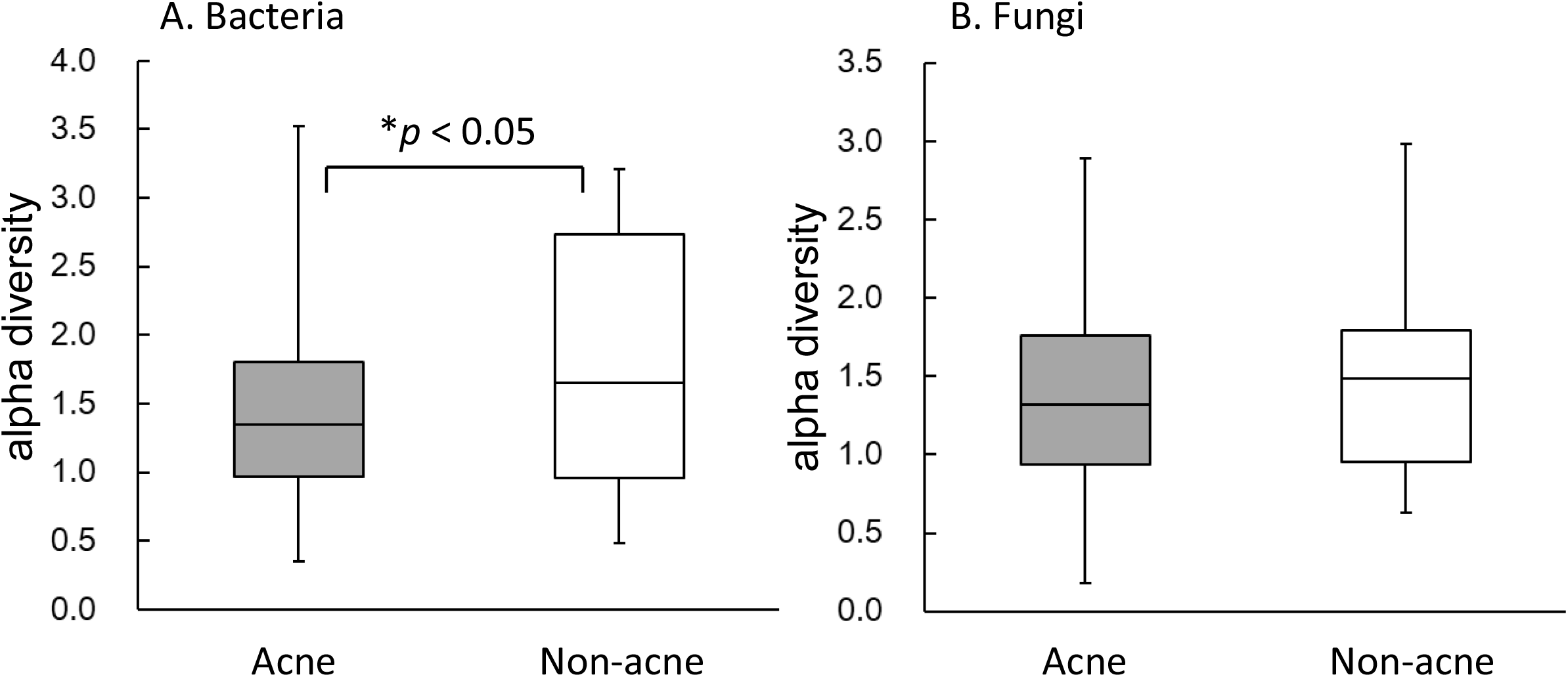
Alpha diversity (Shannon index) of facial skin bacteria (A) or fungi (B) microbiome samples from Acne (n = 56) or Non-acne (n = 20) subjects. The lines inside the rectangles represent the median, and the whiskers indicate the maximum and minimum values. Statistical differences were calculated using Welch’s *t*-test with Holm’s correction.

### Relationship between acne vulgaris and C. acnes

Among the 296 participants, 219 were categorized into the Acne group, identified as having acne vulgaris, while 77 were categorized into the Non-acne group, determined as not having acne. Using the DNA samples extracted from facial skin, the proportion of *C. acnes* within the entire bacterial community was investigated. As a result, the proportion of *C. acnes* was significantly higher in the Acne group (51.7%) compared to the Non-acne participants (44.3%) (*p* < 0.05, Fig. 4-A). The logarithmic count of *C. acnes* per single sticker strip was significantly reduced in the Acne group (4.12) compared to the Non-acne group (3.72) (*p* < 0.001, Fig. 4-B). Furthermore, the proportion of *C. acnes* within all bacteria decreased with age, and the proportion in individuals in their 40s was significantly lower compared to those in their 20s (*p* < 0.05, Fig. 5).

**Figure 4.**
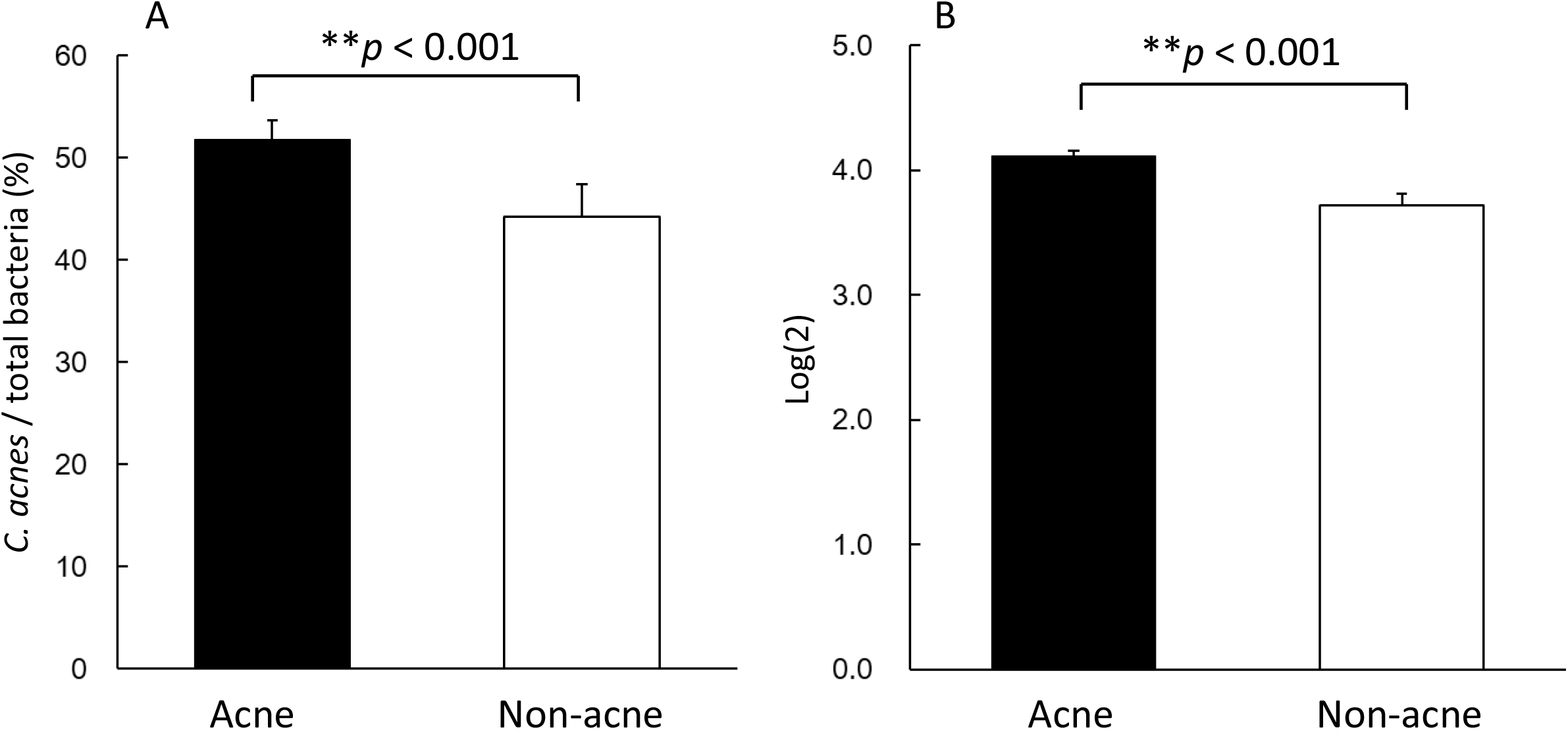
The percentage (A) and log copy number (B) of *C. acnes* among total bacteria in facial skin samples from Acne (n = 219) or Non-acne (n = 77) subjects. Statistical differences were calculated using Welch’s *t*-test with Holm’s correction.

**Figure 5.**
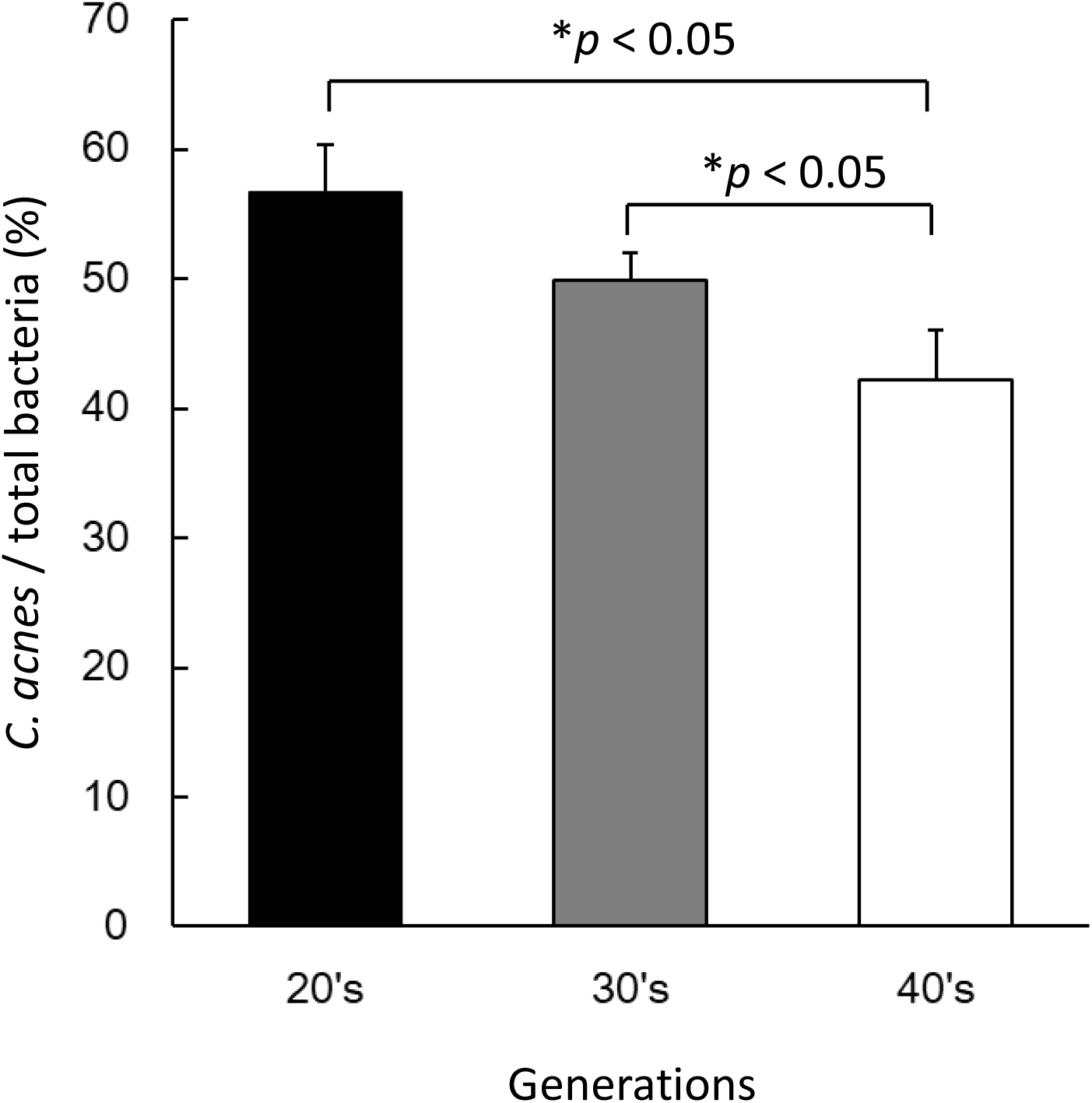
Changes in the proportion of *C. acnes* in total bacteria in facial skin samples were observed among the following age groups: 20s (n = 43), 30s (n = 181) and 40s (n = 70). Statistical differences were calculated using one-way ANOVA followed by the Holm’s *post hoc* test.

### C. acnes phylotypes

Next, we investigated the proportion of *C. acnes* subtypes in all 76 participants. Subtype IA_1_ (42.8%) was the most common, followed by the subtype IA_2_ (24.5%), Type II (21.5%), subtype IB (10.0%), and type III (1.1%), as shown in Fig. 6. When comparing the proportions of *C. acnes* subtypes between the Acne group and the Non-acne group, there was no difference observed in the proportions of IA_1_, IA_2_, Type II, and IB types (data not shown). On the other hand, for Type III, while the Acne group had a proportion of 0.6%, the Non-acne group showed a higher proportion of 2.5%. The Acne group had a significantly (*p* < 0.01) lower proportion of type III as shown in Fig. 7.

**Figure 6.**
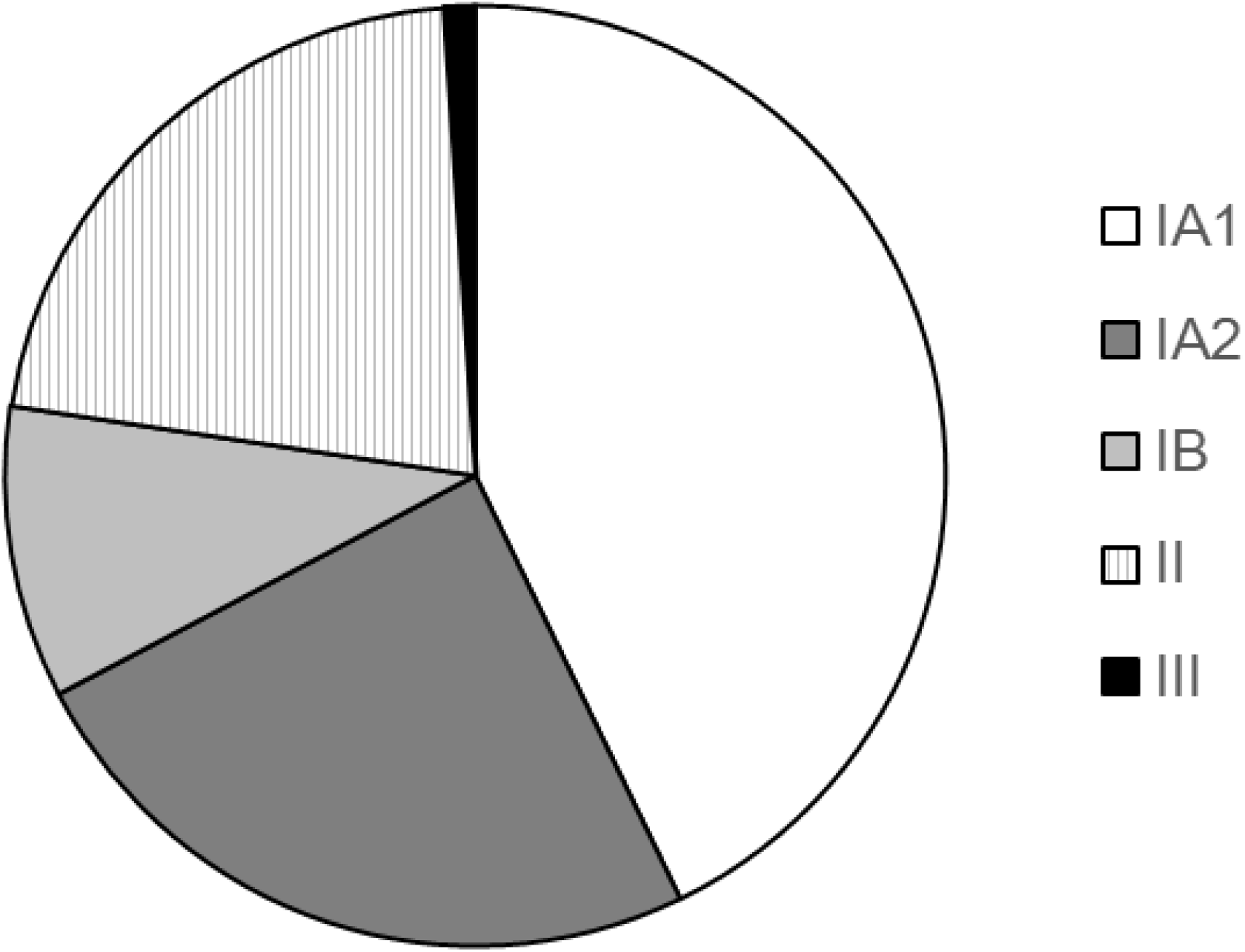
*C. acnes* phylotype composition of all subjects.

**Figure 7.**
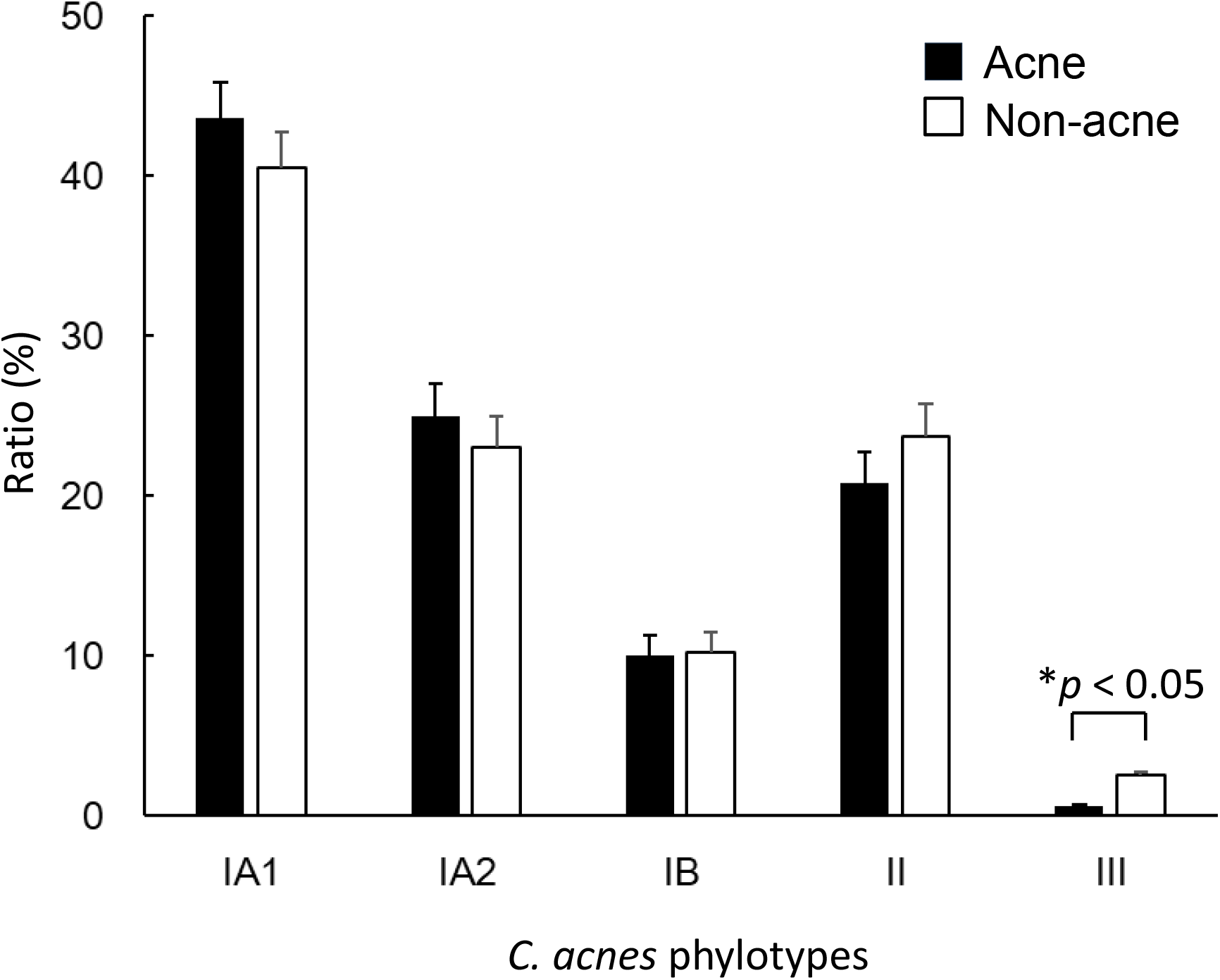
The percentage comparison among *C. acnes* phylotype in facial skin samples from Acne (n = 56) or Non-acne (n = 20) subjects. Statistical differences were calculated using Welch’s *t*-test with Holm’s correction.

### Clonal complex of C. acnes

Upon further investigation of the detailed sequence types (ST) of *C. acnes*, the constituent ratios, starting from the highest, were as follows: IA2_2_F0 23.9%, IA1_4_A0 20.6%, II_2_K0 18.6%, IA1_5_A2 11.1%, IB_1_H0 10.5%, IA1_6_D0 3.0%, IA1_5_A1 3.0%, II_3_K1 2.5%, III_1_L0 2.0%, IA2_1_F1 1.7% (Table 2). The result was sorted by presence or absence of acne vulgaris. As a result, for ST III_1_L0, the Acne group had 2.5% while the Non-acne group had 0.6%, showing a significant difference (*p*<0.01, without correction). For ST IA2_1_F1, the Acne group had 2.3%, while the Non-acne group had 0.2%, displaying a significant difference (*p* < 0.05, without correction) (Table 3). Upon comparison by age group, the biggest differences between age groups (20s and 40s) were found in IA1_4_A0 (11.1%), IA1_5_A2 (6.6%), IA1_6_D0 (−6.0%), and IA2_2_F0 (−6.3%). However, no statistically significant differences were observed.

**Table 2.** The composition of *C. acnes* sequence types (ST)

| Sequence types (ST) | Ratio (%) |
| --- | --- |
| IA2_2_F0 | 23.9 |
| IA1_4_A0 | 20.6 |
| II_2_K0 | 18.6 |
| IA1_5_A2 | 11.1 |
| IB_1_H0 | 10.5 |
| IA1_6_D0 | 3.0 |
| IA1_5_A1 | 3.0 |
| II_3_K1 | 2.5 |
| III_1_L0 | 2.0 |
| IA2_1_F1 | 1.7 |

**Table 3.**
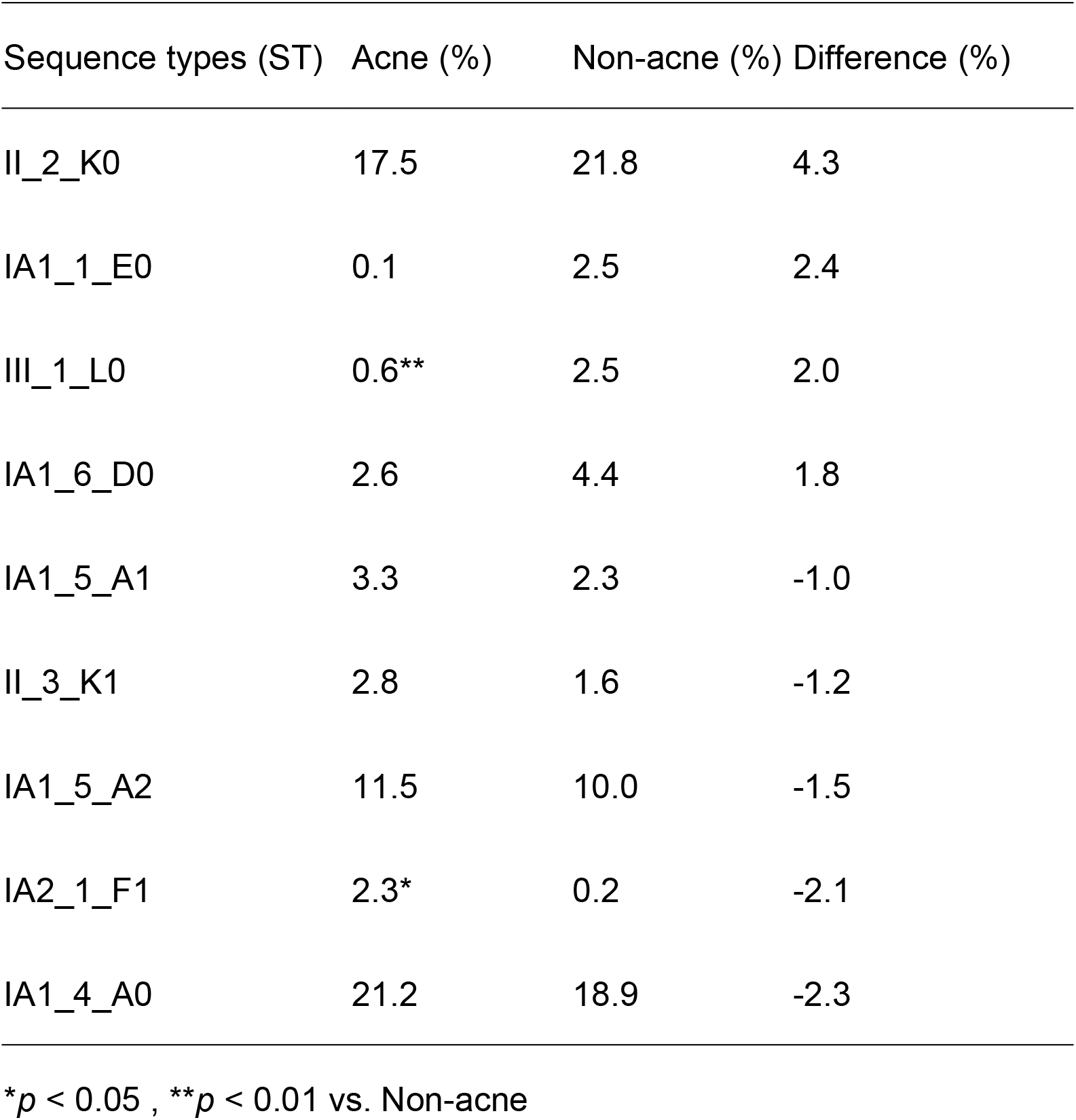
The composition of *C. acnes* sequence types (ST)

## Discussion

There is a growing demand for skin microbiome analysis from the perspective of skin disease treatment and beauty. Consequently, a technique that can sample skin bacteria easily and accurately is highly sought after. Various sampling methods, such as cotton swabs, scrapings, cyanoacrylate gel biopsy, and needle biopsy, have been devised and used in research for collecting the skin microbiome (flora)^20^. Among them, the swabbing method using a moistened cotton swab has been reported to be able to collect a bacterial flora equivalent to that of a biopsy, and it has been relatively commonly used^23,24^. However, the swab method had drawbacks, such as difficulty controlling efficiency depending on the sampling site and technique, and poor preservation.

In this study, we used a patch adhesive sticker with high preservation and adhesiveness (MySkin® patch), allowing subjects to sample microbes from their own facial surface. We evaluated samples sent by post and confirmed that there was no issue using them for microbiome analysis. Previously, Ogai *et al.* collected samples using either the cotton swab method or the sticker-stripping method and investigated the microbial composition using both next-generation sequencing and culture methods. They reported that the results of both methods were equivalent^24^. MySkin® patch is designed for even more convenient collection, such as having a constant seal area, and also possesses excellent bacterial preservation properties, making it possible for subjects to collect samples themselves. Indeed, this study demonstrated that samples could be easily collected and sent by mail.

*C. acnes* is the most abundant resident bacterium of facial skin and is found in both acne patients and healthy subjects^30,31^. Since it is anaerobic, its main niche is believed to be the deeper area of the sebaceous hair follicles^32,33^. Thus, to study the relationship between acne vulgaris and *C. acnes*, methods involving follicle or skin biopsy have been primarily used^20^. However, follicle tissue sampling such as skin biopsy is more challenging. One study that have employed swab sampling, *C. acnes* has been reported to constitute more than 30% of the facial microbial community in acne patients^21^. On the other hand, another report that sampled 55 facial acne patients using the swab method and found that even in acne-affected areas, the proportion of propionic acid-producing bacteria such as *C. acnes* was much lower (less than 2%)^22^.

In this study, we obtained web questionnaires and image analysis from photographs taken with smartphones from subjects to differentiate between the Acne group and the Non-acne group. It is widely known that the proliferation of *C. acnes* is a factor in the onset of acne vulgaris^5,13,34^. In this study, we also examined the relative abundance of *C. acnes* in the samples. As a result, the rate of *C. acnes* was significantly higher in the Acne group. From this, it was believed that even with data from web questionnaires and images taken with smartphones^35,36^, one can grasp the skin condition of the subjects, and it is possible to differentiate between the Acne group and the Non-acne group.

Results of hierarchical clustering analysis on skin resident bacteria showed a group where the microbial community was dominated by *C. acnes*, and the proportion of *C. acnes* was high. When comparing the Acne group and the Non-acne group, both the proportion and copy number of *C. acnes* were significantly higher in the Acne group. However, considering that many subjects were diagnosed with acne vulgaris even in groups dominated by *Neisseriaceae* and others, it is unlikely that *C. acnes* alone has a pronounced influence on the disease. Our new finding is that there might be various factors other than *C. acnes* in the pathogenesis of acne vulgaris. Moreover, the alpha diversity (species richness) of the facial skin microbial community was significantly lower in the Acne group (Fig. 6-A). This result is consistent with the observation that the incidence of acne vulgaris decreases with age, and α-diversity increases^37,38^.

*C. acnes* is generally composed of several phylogenetic subtypes, commonly referred to as IA_1_, IA_2_, IB, IC, II, III^17^. At the genome level, even finer classifications are possible^17,39^. In this study, a genetic analysis was conducted to determine the phylotype ratio of *C. acnes* in all subjects. IA_1_ was found to be the most prevalent type, consistent with a previous report^40^. It has been reported that the IA_1_ type, isolated from common acne, possesses strong pathogenicity^13,18,19^. In the hair follicles of acne patients, the dominant strain of *C. acnes* is IA_1_. On the surface of the face, not only IA_1_ but also IA_2_, IB, and II are present^40,41^. When we examined the phylotype of *C. acnes* on the facial skin surface, we found a cocktail of IA_1_, IA_2_, IB, II, and III, with no significant difference between the Acne and Non-acne groups. This result may be due to the wider area of sampling rather than pinpoint acne lesions. Meanwhile, type III, which showed a significant difference between the groups, has been found in normal skin but not in common acne areas^14,19^. In our research, type III was confirmed in both Acne and Non-acne groups. It’s suggested that the reason that type III, which is rarely reported in the face, was found in the Acne group might be due to the inability to strictly separate common acne lesion from surrounding areas during sticker sampling. Further detailed verification is needed in the future, but for narrow lesion sampling, biopsies or swabs might be suitable. Type III is reported to lack toxic factors and has a different morphology and metabolic pathway than types I and II^42^. In our results, type III in the Acne group showed significantly lower values compared to the Non-acne group. Further analysis is needed to understand how the decreased expression of type III in normal skin areas relates to the onset of common acne.

When examining the detailed sequence types of *C. acnes*, the strains with the highest proportion in all subjects were IA2_2_F0 (23.9%), IA1_4_A0 (20.6%), and II_2_K0 (18.6%). IA2_1_F1 (*p* < 0.05 vs Non-acne group) higher in acne patients, while III_1_L0 was lower (*p* < 0.01 vs Non-acne group) (Table 3, Supplemental figure 2). Also, our study revealed that various types of *C. acnes* phylotypes were expressed on the skin surface of the face. Our method of using specialized sticker to collect samples from specific facial areas captures both acne and healthy tissues. This may indicate skin conditions prone to developing acne. Dagnelie *et al*. reported that the detailed phylotype expression patterns of *C. acnes* differ with the onset of acne^40^. In the future, increasing the number of subjects and conducting larger-scale analyses may clarify statistical differences between groups.

In this study, we could successfully conduct postally collected skin microbiome samples and made analysis by online basis from customers of a company. Although preliminary, our results suggest the potential for online consultations to provide diagnostic quality comparable to in-person assessments in testing and diagnosing skin-related diseases. Especially in the post-COVID era, the advancement of such research could significantly contribute to the diversification of diagnostic methods and the development of dermatology.

