## Supplementary material for "Detailed microbiome analysis of sticker-stripped surface materials of acne lesions revealed acne-related *Cutibacterium acnes* subtypes: a pilot study": Suppl text

New Results

1). KINS LAB, Tokyo, Japan

2). MySkin Corporation, Tokyo, Japan

3). Department of Dermatology, The Jikei University School of Medicine, Tokyo, Japan

*Correspondence: Itaru Dekio, M.D., Ph.D.

Department of Dermatology, School of Medicine, The Jikei University School of Medicine

3-25-8 Nishi-shinbashi, Minato-ku, Tokyo 105-8461, Japan

**Figure Legends**

Supplemental figure 1

Composition of *C. acnes* sequence types (ST) in all subjects (n = 76).

Supplemental figure 2

*C. acnes* sequence types (ST) composition between acne (n = 56) and non-acne (n = 20) subjects.
