## Supplementary figures and images for "Detailed microbiome analysis of sticker-stripped surface materials of acne lesions revealed acne-related *Cutibacterium acnes* subtypes: a pilot study"

### Suppl Figures

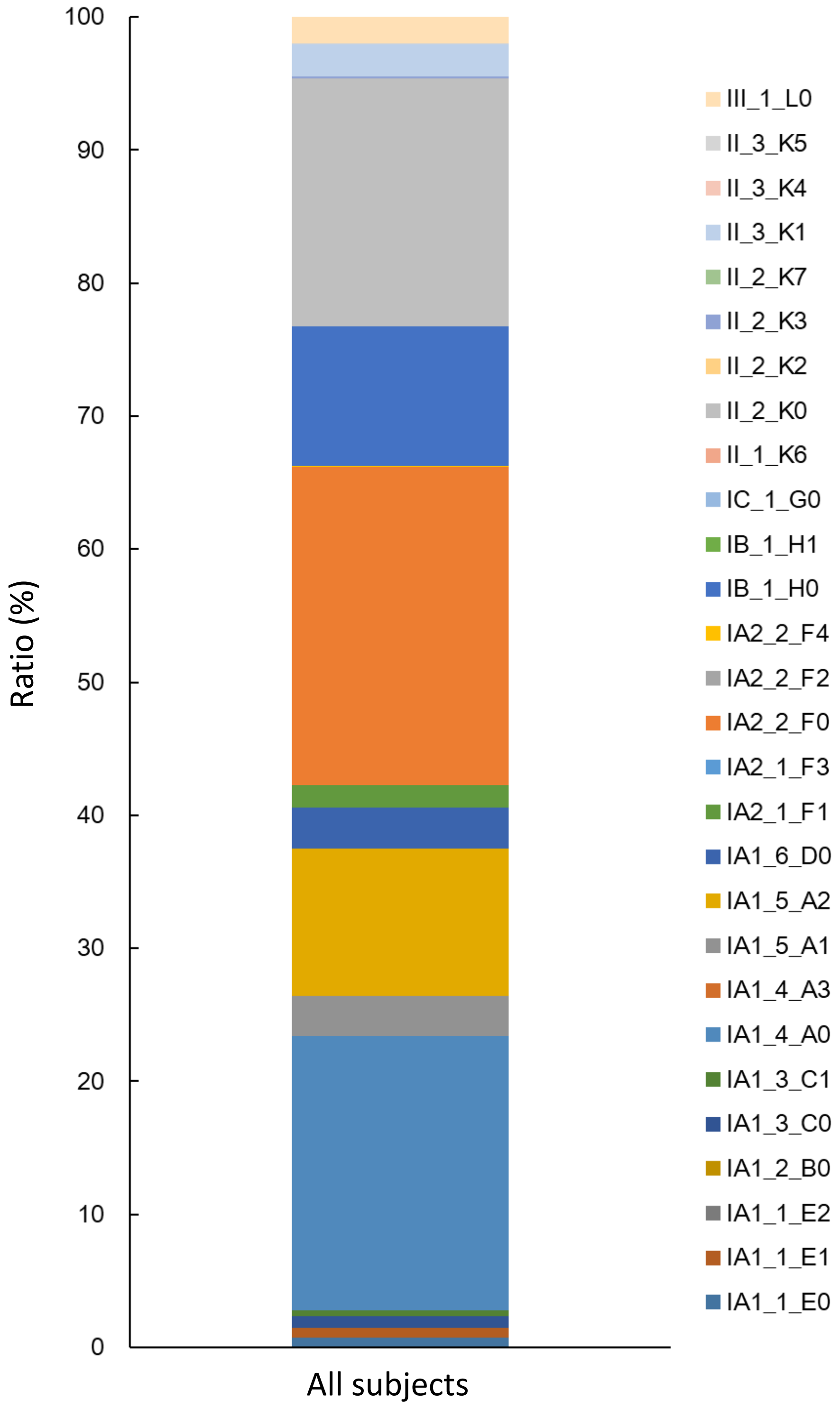

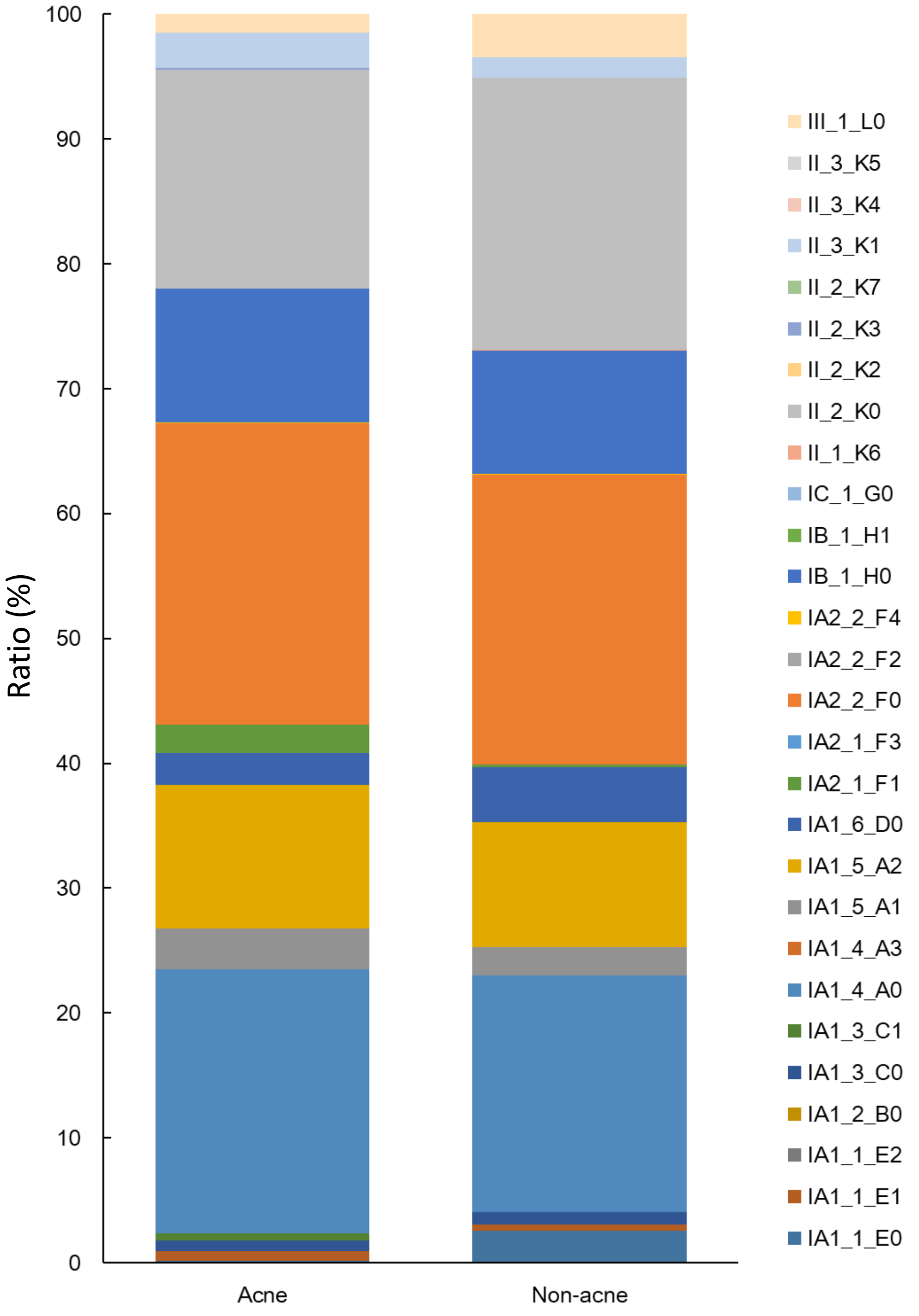
